# Ventricular Expansion Couples Hyperosmotic Stress to Thirst

**DOI:** 10.64898/2026.06.13.732103

**Authors:** Tingting Zhu, Chaohua Jiang, Jin Yang, Hanming Zheng, Xiaoli Lin, Weizhen He, Lijie Zhang, Zihao Chen, Yun Stone Shi, Haiyun Ren, Zhihai Qiu

**Author notes:** Corresponding authors: Zhihai Qiu, Ph.D. Haiyun Ren, Ph.D. These authors contributed equally to this work (Tingting Zhu, Chaohua Jiang).

## Abstract

Thirst is thought to arise from osmotic signals detected by circumventricular organs, yet how hyperosmotic stress activates mechanosensitive channels remains unclear because cell shrinkage should reduce membrane tension. Here we identify a brain-scale mechanical signal that couples systemic osmotic stress to drinking. In mice, hypertonic saline rapidly increased serum and cerebrospinal fluid osmolality, expanded the lateral ventricles, and promoted water intake. Relieving or blocking ventricular deformation attenuated drinking without eliminating osmotic gradients. Hyperosmotic challenge also produced localized deformation of periventricular cells, where spatial transcriptomics revealed candidate mechanosensitive channels, including Tmem63b and Piezo1. Thus, ventricular expansion provides a mechanical component of osmotic thirst, uncovering an osmo-mechanical layer of interoceptive regulation.

## Introduction

Maintenance of body fluid homeostasis is essential for survival. Increases in plasma osmolality dehydrate cells and elicit thirst (*1*), a powerful interoceptive drive that promotes water seeking and drinking. This response is classically attributed to osmosensory neurons in the lamina terminalis, including the subfornical organ (SFO) and organum vasculosum of the lamina terminalis (OVLT), which lack a conventional blood–brain barrier and directly monitor circulating osmotic signals (*2-4*). Together with the median preoptic nucleus, these structures form a forebrain network that converts hyperosmotic stress into neural activity and motivated drinking (*2, 4, 5*).

A major unresolved question is how osmotic stress is converted into the neural signals that initiate thirst. Mechanosensitive ion channels have long been implicated in osmosensation because they can couple membrane deformation to ionic flux and neuronal excitability (*6*). Several channel families, including Trpv channels and the Tmem63 family, have been linked to osmotic responses in sensory and central neurons (*7-11*). Recent work has further suggested that Tmem63b functions as a hyperosmolality sensor in SFO neurons (*10, 11*), supporting a role for mechanically gated or mechanically tuned channels in osmotic thirst. However, this model presents a biophysical paradox. Most mechanically activated channels are favored by increased membrane tension (*6*), as occurs during hypoosmotic swelling, whereas hyperosmotic stress drives water out of cells, causing shrinkage and reducing global membrane tension. How, then, are mechanosensitive pathways engaged during hyperosmotic thirst?

One possibility is that the relevant mechanical signal does not arise solely at the single-cell membrane. The brain is a fluid-filled organ in which osmotic stress can alter water distribution among cells, interstitial space, blood, and cerebrospinal fluid (*12*). Because osmosensory nuclei are positioned adjacent to the ventricular system (*3, 5*), osmotic shifts could reshape local tissue architecture and impose mechanical forces on periventricular cells. Such tissue-scale mechanics have not been incorporated into current models of thirst, which have mainly emphasized molecular osmosensors and neural circuits (*4, 5, 13, 14*).

Here we identify ventricular dynamics as a mechanical component of osmotic thirst. Using functional ultrasound imaging in mice challenged with hypertonic saline, we found that systemic hyperosmotic stress rapidly enlarged the lateral ventricles while increasing serum and cerebrospinal fluid osmolality and inducing robust drinking. Releasing ventricular fluid through an open cannula reduced hypertonicity-induced ventricular enlargement and suppressed water licking, indicating that ventricular expansion contributes to the behavioral expression of thirst rather than simply accompanying osmotic stress. Histological analysis further revealed localized deformation of nuclei adjacent to the ventricular wall, and spatial transcriptomic profiling identified broad periventricular expression of candidate mechanosensitive channels, particularly Tmem63b and Piezo1. These findings suggest that hyperosmotic stress generates a brain-scale mechanical signal through ventricular expansion, providing a force source for periventricular mechanosensation and defining an osmo-mechanical layer of interoceptive regulation.

## Results

### Hyperosmotic challenge rapidly expands the lateral ventricles

To determine how systemic hyperosmotic stress alters brain physiology during thirst, we established an osmotic thirst model by intraperitoneal injection of hypertonic NaCl. Mice received either 2 M NaCl or isotonic 0.15 M NaCl and were water-deprived for 0, 10, 20, 30, 40, 50, or 60 min before a 10-min water intake test (Fig. 1A). Hypertonic NaCl rapidly increased water licking, which peaked at 20 min and then gradually declined despite continued water deprivation (Fig. 1B). Thus, osmotic drinking was rapidly induced but did not increase linearly with deprivation time.

**Fig. 1.**
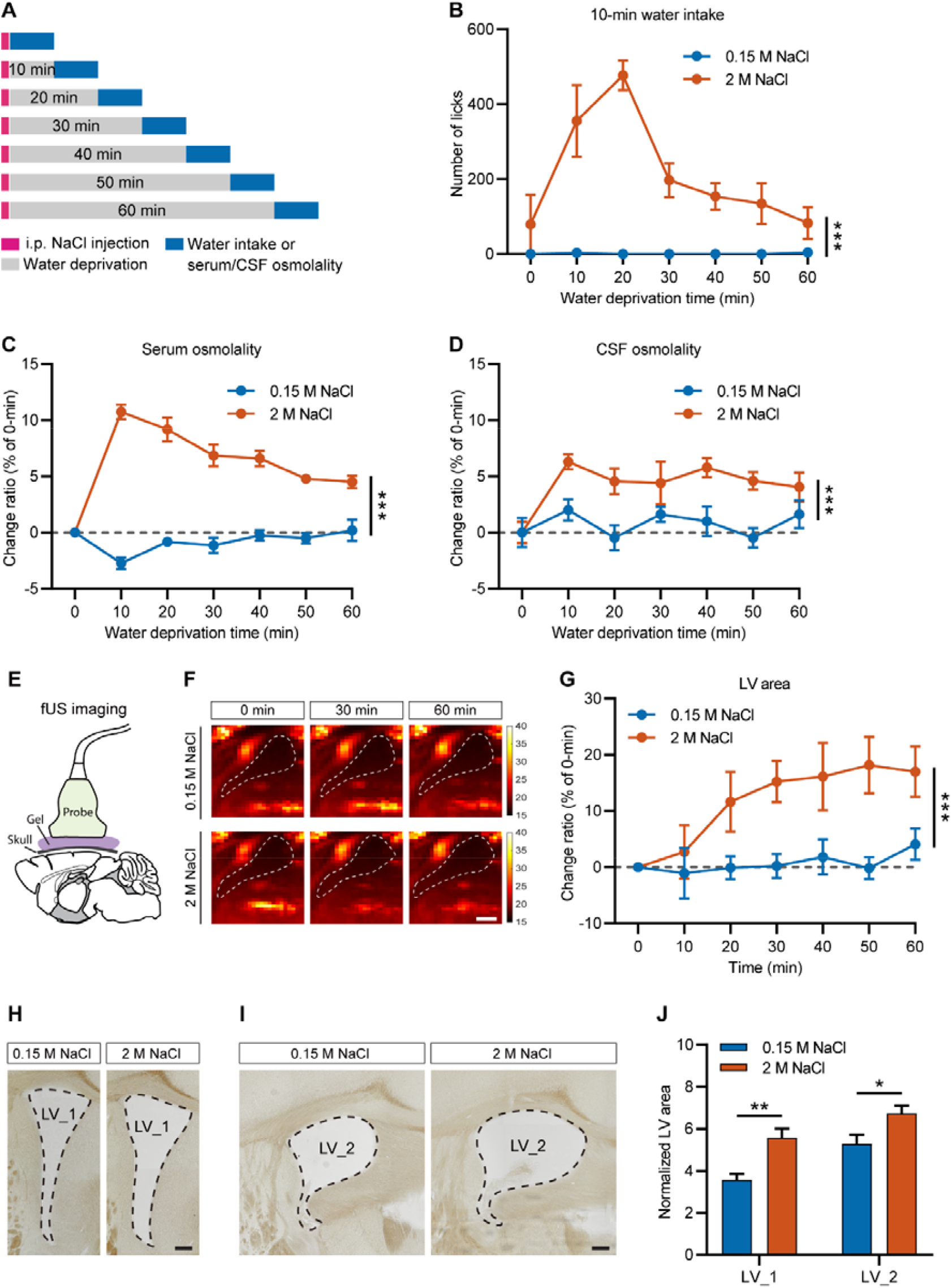
Hyperosmotic challenge rapidly expands the lateral ventricles. (**A**) The diagram shows experimental procedures of water intake, serum, and CSF osmolality measurement after indicated time of i.p. injection of NaCl. (**B**) Number of water licks for 10 minutes at indicated time after i.p. injection of NaCl (n = 5-8 mice). (**C**) The change ratio of serum osmolality at indicated time after i.p. injection of NaCl (0.15 M NaCl: n = 5 mice; 2 M NaCl: n = 8 mice). (**D**) The change ratio of CSF osmolality at indicated time after i.p. injection of NaCl (0.15 M NaCl: n = 4 mice; 2 M NaCl: n = 6 mice). (**E**) The diagram shows fUS imaging of mouse brain. (**F**) Representative fUS images under sagittal view of the brain collected at 0, 30, and 60 min, respectively, after i.p. injection of NaCl; the lateral ventricle (LV) area is marked by the white dotted line; scale bar: 500 µm. (**G**) The change ratio of LV area at indicated time after i.p. injection of NaCl (0.15 M NaCl: n = 6 mice; 2 M NaCl: n = 7 mice). (**H**) Representative images of the LV area (∼ 0.1 mm relative to Bregma) under isotonic (0.15 M NaCl) or hypertonic (2 M NaCl) condition; the LV area is marked by the black dotted line; scale bar: 200 µm. (**I**) Representative images of the LV area (∼ -0.5 mm relative to Bregma) under isotonic (0.15 M NaCl) or hypertonic (2 M NaCl) condition; the LV area is marked by the black dotted line; scale bar: 200 µm. (**J**) Statistical analysis of the LV area (0.15 M NaCl: n = 7 mice for both LV_1 and LV_2; 2 M NaCl: n = 8 mice for both LV_1 and LV_2). All data were presented as mean ± SEM. Significant differences between isotonic (0.15 M NaCl) and hypertonic (2 M NaCl) groups were analyzed by two-way ANOVA and denoted as * (p < 0.05), ** (p < 0.01), or *** (p < 0.001).

We next asked whether this nonlinear behavioral response reflected changes in body fluid osmolality. Serum or cerebrospinal fluid (CSF) was collected at the same time points after NaCl injection (Fig. 1A). In mice challenged with 2 M NaCl, both serum and CSF osmolality increased rapidly and remained elevated over the observation period, whereas isotonic NaCl produced little change (Fig. 1C and D). These results indicate that systemic hyperosmotic challenge rapidly alters both peripheral and central osmotic environments.

To examine brain-wide physiological changes associated with this osmotic state, we performed functional ultrasound imaging (Fig. 1E). Unexpectedly, hypertonic NaCl induced a marked enlargement of the lateral ventricle area over time (Fig. 1F and G). This effect was confirmed histologically: brain sections prepared under matched osmotic conditions showed larger lateral ventricles in hypertonic NaCl-treated mice than in isotonic controls at two anatomical levels (Fig. 1H to J). Thus, systemic hyperosmotic challenge induces rapid and sustained ventricular expansion.

### Ventricular fluid release attenuates hypertonicity-induced ventricular expansion and drinking

Hyperosmotic stress draws water out of cells, causing cellular shrinkage and redistribution of fluid across brain compartments. We therefore asked whether ventricular fluid accumulation contributes to the observed ventricular expansion. To test this, we implanted a cannula into the lateral ventricle and opened it during hypertonic challenge to release ventricular fluid (Fig. 2A).

**Fig. 2.**
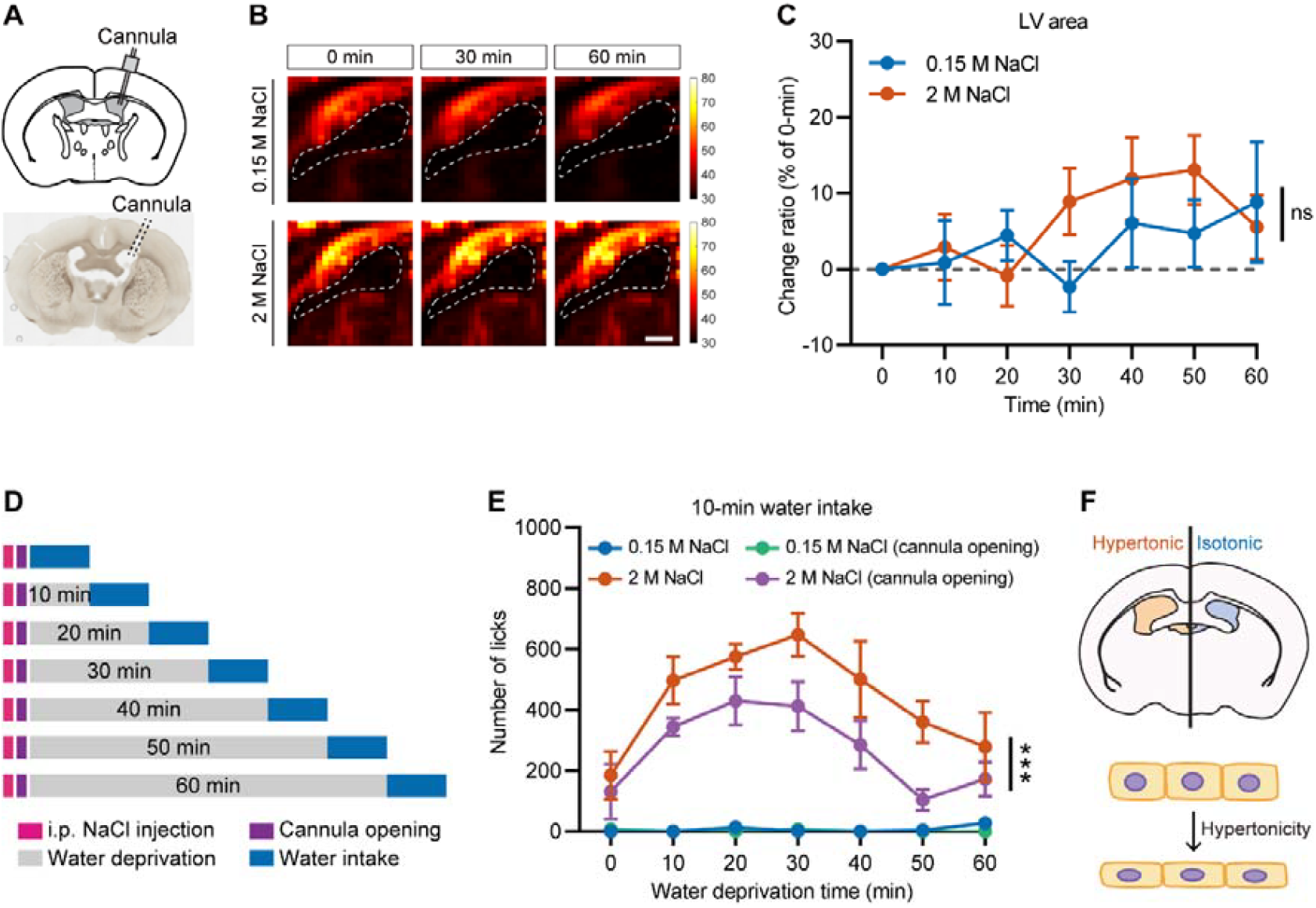
Release of the excessive CSF suppresses hypertonicity-induced LV enlargement and thirst. (**A**) The diagram (top) and image (bottom) show implantation of a cannula into the LV. (**B**) Representative fUS images under sagittal view of the brain collected at 0, 30, and 60 min, respectively, after i.p. injection of NaCl and CSF release through the cannula; the lateral ventricle (LV) area is marked by the white dotted line; scale bar: 500 µm. (**C**) The change ratio of LV area at indicated time after i.p. injection of NaCl and CSF release (0.15 M NaCl: n = 4 mice; 2 M NaCl: n = 7 mice). (**D**) The diagram shows experimental procedures of water intake after indicated time of i.p. injection of NaCl with or without CSF release through the cannula. (**E**) Number of water licks for 10 minutes at indicated time after i.p. injection of NaCl with or without CSF release through the cannula (n = 6-16 mice). (**F**) The diagram shows that hypertonic challenge induces deformation of the LV and periventricular cells. All data were presented as mean ± SEM. Significant differences between isotonic (0.15 M NaCl) and hypertonic (2 M NaCl) groups were analyzed by two-way ANOVA and denoted as ns (not significant) or *** (p < 0.001).

With the ventricular cannula open, hypertonic NaCl no longer produced a significant increase in lateral ventricle area during fUS imaging (Fig. 2B and C). This intervention also altered behavior. Mice with an open ventricular cannula showed reduced water licking after 2 M NaCl injection compared with hypertonic controls without ventricular release (Fig. 2D and E). These findings indicate that ventricular expansion is not merely a passive anatomical consequence of hyperosmotic stress, but contributes to the full behavioral expression of osmotic thirst.

### Hyperosmotic ventricular expansion deforms periventricular cells

We next asked whether ventricular expansion could generate local mechanical deformation in surrounding brain tissue. Hyperosmotic stress may influence periventricular cells through two related processes: cell shrinkage caused by intracellular dehydration and tissue deformation caused by ventricular enlargement (Fig. 2F). We therefore examined nuclear morphology as a cellular readout of local mechanical deformation.

Mice were injected with isotonic or hypertonic NaCl, and brains were fixed and sectioned under corresponding osmotic conditions (Fig. 3A). We focused on three regions of interest around the lateral ventricle, including dorsal, septal, and ventral periventricular regions, and subdivided each region into ten layers according to distance from the ventricular wall (Fig. 3A). In the dorsal periventricular region, hypertonic challenge significantly increased the nuclear aspect ratio in the first layer adjacent to the ventricle (Fig. 3B and C). In contrast, deeper layers and the other two regions showed no evident change in nuclear aspect ratio (Fig. S1A, B, D, and E). Interestingly, all three regions exhibited decreased nuclear area in response to hypertonic challenge (Fig. 3D, S1C, and S1F). These data suggest that hyperosmotic ventricular expansion produces spatially restricted cellular deformation near the ventricular wall, rather than uniform tissue deformation throughout the periventricular area.

**Fig. 3.**
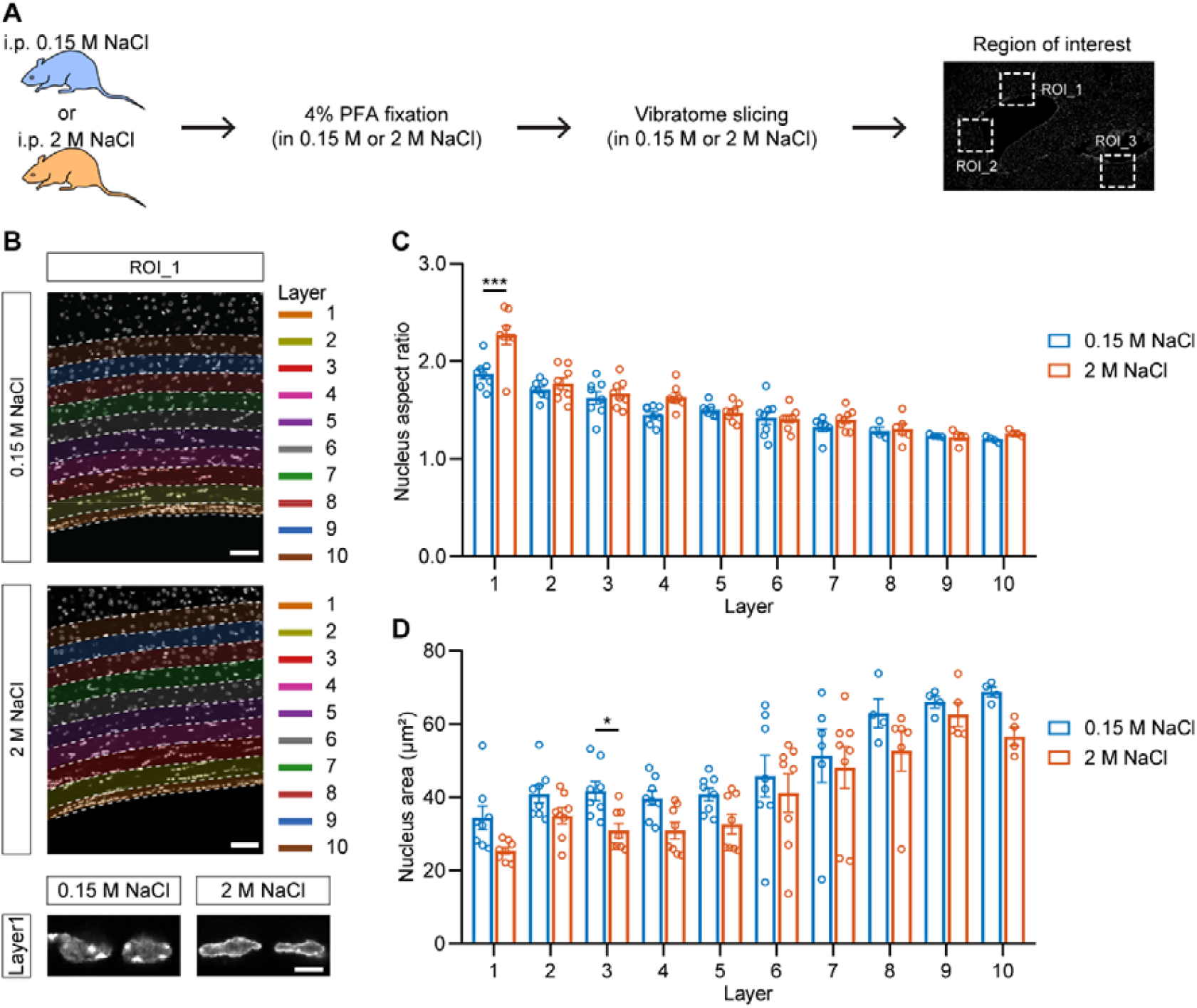
Hyperosmotic ventricular expansion deforms periventricular cells. (**A**) The diagrams show experimental procedures of brain sections prepared under isotonic (0.15 M NaCl) or hypertonic (2 M NaCl) condition; the three regions of interest are marked with white dotted frames. (**B**) Representative images of the ten layers of nuclei (top and middle, scale bar: 50 µm) and single nucleus in layer 1 (bottom, scale bar: 5 µm) of ROI_1 under isotonic (0.15 M NaCl) or hypertonic (2 M NaCl) condition. (**C**) Statistical analysis of nucleus aspect ratio of cells in each layer under isotonic (0.15 M NaCl) or hypertonic (2 M NaCl) condition (n = 4-8 mice). (**D**) Statistical analysis of nucleus area of cells in each layer under isotonic (0.15 M NaCl) or hypertonic (2 M NaCl) condition (n = 4-8 mice). All data were presented as mean ± SEM. Significant differences between isotonic (0.15 M NaCl) and hypertonic (2 M NaCl) groups were analyzed by two-way ANOVA and denoted as * (p < 0.05) or *** (p < 0.001).

### Periventricular cells express candidate mechanosensitive channels

Because ventricular expansion produced local deformation of periventricular cells, we next asked whether these cells express mechanosensitive ion channels that could detect such mechanical changes. We performed MERFISH to map the expression of candidate mechanosensitive channels around the lateral ventricle. Two periventricular subregions were selected, and each was divided into ten layers from the ventricular wall outward (Fig. 4A).

**Fig. 4.**
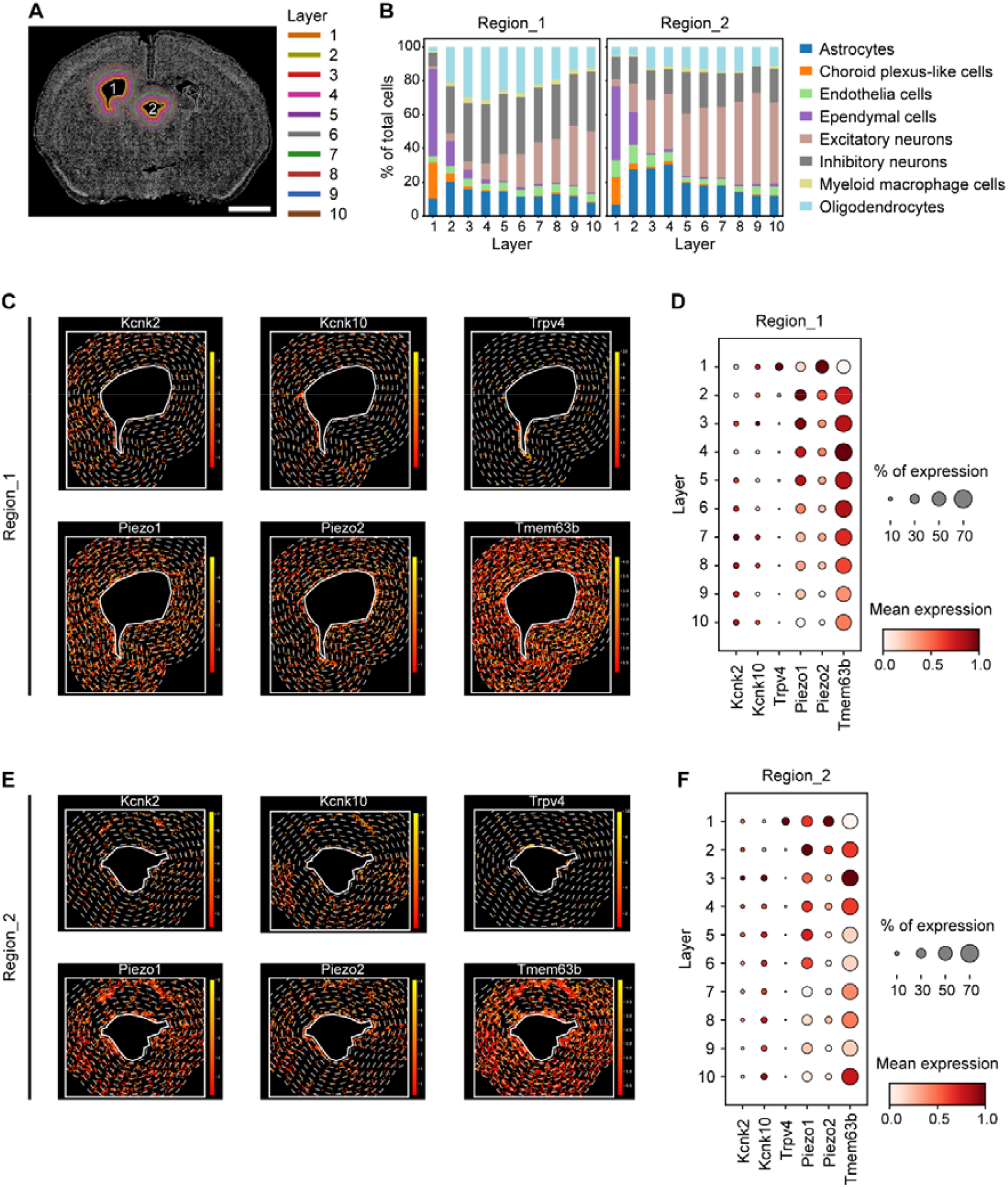
Spatial expression of candidate mechanosensitive ion channels in periventricular cells. (**A**) The representative image shows ten layers of cells in the two ventricular regions; scale bar: 1 mm. (**B**) Percentage of different types of cells in each layer within each ventricular region. (**C**) The representative images show spatial expression of Kcnk2, Kcnk10, Trpv4, Piezo1, Piezo2, and Tmem63b in region_1, each of the ten layers is separated by white dotted lines. (**D**) The bubble plots show the mean and percentage expression of Kcnk2, Kcnk10, Trpv4, Piezo1, Piezo2, and Tmem63b in each layer within region_1. (**E**) The representative images show spatial expression of Kcnk2, Kcnk10, Trpv4, Piezo1, Piezo2, and Tmem63b in region_2, each of the ten layers is separated by white dotted lines. (**F**) The bubble plots show the mean and percentage expression of Kcnk2, Kcnk10, Trpv4, Piezo1, Piezo2, and Tmem63b in each layer within region_2.

Cell-type analysis showed that both subregions contained neurons and non-neuronal cells, with broadly similar cellular composition but different cell numbers across layers (Fig. 4B). We then analyzed the layer-by-layer expression of six candidate mechanosensitive channels: Kcnk2, Kcnk10, Trpv4, Piezo1, Piezo2, and Tmem63b. Among these, Tmem63b and Piezo1 were broadly expressed across both periventricular subregions and across multiple layers (Fig. 4C to F). Piezo2 showed more moderate expression, whereas Kcnk2, Kcnk10, and Trpv4 were more sparsely distributed (Fig. 4C to F).

Together, these results identify Tmem63b and Piezo1 as prominent candidate mechanosensitive channels in the periventricular tissue surrounding the lateral ventricle. Combined with the nuclear deformation data, these findings support a model in which hyperosmotic stress expands the ventricular system, mechanically deforms nearby cells, and recruits periventricular mechanosensitive pathways to modulate osmotic thirst.

## Discussion

Homeostatic regulation of body fluid osmolality is fundamental for survival, because even small deviations in extracellular osmolality can impose molecular, cellular, and systemic stress (*1*). In this study, we identify a previously unappreciated brain-scale response to hyperosmotic challenge: systemic hypertonicity rapidly enlarges the lateral ventricles and deforms periventricular cells. In mice challenged with intraperitoneal 2 M NaCl, serum and cerebrospinal fluid (CSF) osmolality increased rapidly, accompanied by robust osmotic drinking. Unexpectedly, functional ultrasound imaging and histological validation revealed a significant enlargement of the lateral ventricles under hypertonic conditions. Releasing ventricular fluid through an open cannula attenuated ventricular enlargement and reduced water licking, suggesting that ventricular expansion is not merely a passive anatomical consequence of hypertonicity, but contributes to the behavioral expression of osmotic thirst. At the cellular level, hypertonicity produced a spatially restricted increase in nuclear aspect ratio in the cell layer adjacent to the ventricular wall, indicating local deformation of periventricular cells. Spatial transcriptomic analysis further showed broad periventricular expression of candidate mechanosensitive channels, particularly Tmem63b and Piezo1. Together, these findings support a model in which hyperosmotic stress generates not only molecular osmotic signals but also a tissue-scale mechanical signal through ventricular expansion, providing a potential force source for mechanosensitive pathways during thirst.

Our findings extend a long history of work on osmotic thirst across species. Classic studies in goats showed that hypertonic NaCl delivered into the third brain ventricle can evoke drinking (*15*), antidiuretic hormone release (*16*), and natriuresis (*17*), establishing the ventricular and periventricular region as a key interface for central control of water balance. In mice, modern circuit and molecular studies have focused mainly on lamina terminalis neurons (*18-23*), especially SFO and OVLT neurons (*7, 19, 20, 22*), and on candidate osmotic sensors such as TRPV-family channels and Tmem63b (*7-11, 20*). Recent works showed that Tmem63b is highly expressed in the SFO and is required for hyperosmolarity-induced drinking (*10, 11*), supporting a mechanosensitive-channel model of osmotic thirst. In humans, a systematic review estimated the thirst threshold near 285 mOsm/kg and found that thirst intensity rises as plasma osmolality increases (*24*). These studies collectively establish that osmotic information can be detected with high sensitivity across species, but they leave a biophysical question unresolved: if hyperosmotic stress causes cells to shrink and reduces global membrane tension, where does the mechanical force come from to engage mechanosensitive ion channels? Our study addresses this gap by shifting attention from single-cell osmosensors alone to the surrounding tissue architecture. We propose that ventricular expansion generated by hyperosmotic stress imposes local deformation on periventricular cells, thereby providing a tissue-level mechanical cue that may cooperate with molecular osmosensors during thirst.

This tissue-level view has broader implications for brain function. The brain is not only an electrical and chemical organ; it is also a soft, fluid-filled mechanical system. Mechanical cues such as tissue deformation, fluid pressure, shear stress, and local stiffness are increasingly recognized as regulators of neural and glial cell behavior (*6, 25*), and mechanosensitive ion channels can convert such forces into electrical or biochemical signals (*6*). However, most studies of thirst and interoception have been framed around two levels: molecular receptors that sense osmotic state and neural circuits that generate motivated behavior (*4, 10, 11, 13, 18-20*). Our results suggest an additional intermediate level: physiological changes can remodel brain tissue geometry, and this remodeling can feed back onto local cells as a functional mechanical signal. In this framework, ventricular expansion is not simply a structural readout of altered osmotic state; it may be an active component of the sensory transformation that links body-fluid imbalance to behavioral output. This concept broadens the traditional view of thirst from a receptor-and-circuit model to a fluid–tissue–cell model, in which osmotic stress, CSF/ventricular dynamics, periventricular deformation, and mechanosensitive pathways act together. More generally, our findings suggest that dynamic changes in brain fluid compartments may regulate neural function under physiological conditions, not only in pathological states such as hydrocephalus (*26*), aging (*27*), or brain injury (*28*). By introducing tissue-scale mechanics into the study of homeostatic behavior, this work opens a new direction for understanding how the brain senses internal bodily states and converts them into motivated actions.

Although our findings identify ventricular expansion as a tissue-scale mechanical component of osmotic thirst, several questions remain unresolved. Ventricular fluid release reduced both ventricular enlargement and drinking, but this manipulation does not directly quantify CSF volume, flow, pressure, or local tissue stress, and therefore the fluid-dynamic origin of ventricular expansion remains to be determined. In addition, periventricular deformation was inferred from fixed-tissue nuclear morphology rather than direct in vivo measurements of tissue strain or membrane tension. The spatial transcriptomic data identify Tmem63b and Piezo1 as candidate mechanosensitive channels, but functional studies using cell-type-specific perturbation, electrophysiology, or calcium imaging are needed to establish their causal roles. Finally, the present work uses an acute mouse hypertonicity model; whether similar ventricular osmo-mechanical coupling occurs during physiological dehydration, chronic osmotic stress, disease states, or across species remains unknown. Future studies integrating live imaging, CSF dynamics, mechanical modeling, and channel-specific manipulations will be required to define how ventricular mechanics are converted into thirst-related neural and behavioral responses.

## Materials and Methods Animals

All animal experiments were performed in accordance with the guidelines of the Institutional Animal Care and Use Committee of Guangdong Institute of Intelligent Science and Technology. We used wild-type C57BL/6 male mice aged between 8-to 12-weeks. Mice were housed in specific pathogen-free (SPF) facilities with a standard 12-h light/dark cycle and freely access to food and water unless otherwise indicated. Isotonicity and hypertonicity was induced by intraperitoneal (i.p.) injection of 0.15 M and 2 M NaCl (5 ul/kg), respectively.

### Water intake

Water intake was measured using an automated lickometer system (Panlab). Mice were singlely housed in the testing chamber for at least one day before the experiment. After i.p. injection of NaCl, mice were deprived water for indicated time before water intake measurement. The number of water licks was recorded for 10 minutes. For CSF release in mice, the cannula was opened right after NaCl injection, followed by water lick measurement.

### Osmolality measurement

After i.p. injection of NaCl, mice were deprived water for indicated time before blood or cerebrospinal fluid (CSF) collection. Briefly, blood was collected from the retro-orbital sinus via a glass tubing by capillary action; plasma was isolated by centrifugation at 1000 rpm for 20 min at 4 □. The CSF was collected from the fourth ventricle via a glass micropipette. The osmolality of plasma and CSF was measured using a vapor pressure osmometer (VAPRO).

### Surgery

For cranial window implantation, mice were anesthetized by i.p. injection of zoleti 50 (50 mg/kg) combined with xylazine hydrochloride (10 mg/kg) and fixed on a stereotaxic apparatus. Hair was shaved and skull was exposed by a sagittal incision along the head. A craniotomy from the anterior to the posterior fontanelle was made with a hand drill. Saline was regularly added during the drilling to minimize the potential thermal damage to the brain. Absorbable gelatin sponges were applied in case of bleeding. Once the bone margins became thin enough, the skull flap was carefully lifted, with the dura mater remained intact. The exposed surface was saturated with saline and a transparent polymethylpentene membrane was placed over it as an acoustic window. The membrane was secured with dental cement. For experiments requiring repeated imaging or probe stabilization, a head plate was implanted with dental cement. After the surgery, mice were daily injected with ceftiofur sodium (20 mg/kg) for 3 days and allowed to recover for 7 days before functional ultrasound imaging.

For CSF release experiments, a cannula was implanted into the lateral ventricle (AP, -0.85 mm; ML, -2.1 mm; DV, -1.45 mm) at a 20° angle relative to the vertical axis. The cannula was secured on the skull with dental cement.

### Multiplexed FISH imaging

Mice were transcardially perfused with RNase-free PBS, the brain was extracted and immediately embedded in OCT, followed by snap-frozen on dry ice and stored at −80°C until sectioning. 10-µm sections were prepared by cryosectioning at −18°C, followed by fixation in 4% PFA for 15 min at room temperature. Transcripts of interest were labeled and imaged using a workflow combining padlock probe hybridization, ligation, rolling circle amplification (RCA), and multiplex FISH. Target probe hybridization and RCA were performed with the Bassfish Sample Preparation Reagent Kit (Bassfish Inc.) according to the manufacturer’s instructions.

Automated fluidics and multi-round imaging were carried out on a MetaScope system (Bassfish Inc.) equipped with a 40× air objective and the Bassfish Sample Image Reagent Kit. Z-stacks were acquired with a 1.5-µm step size across five channels—DAPI (405 nm), AF488 (488 nm), Cy3 (561 nm), Cy5 (640 nm), and Cy7 (750 nm). Each field of view comprised 2048 × 2048 pixels at 0.16 µm per pixel, and adjacent images were collected with 10% overlap to enable downstream stitching. Image deconvolution, maximum intensity projection, stitching, multi-round registration, cell segmentation, transcript spot detection, and gene decoding were performed using the BassAnalyzer software (Bassfish Inc.).

### Functional ultrasound (fUS) imaging

fUS imaging was performed as previously described (*29*), with a linear ultrasound transducer (15 MHz, 110 µm pitch, 128 elements and 8 mm elevation focus) on a Verasonics platform controlled by a modified MATLAB-based miniscan interface. A set of seven tilted plane-wave images (-6, -4, -2, 0, 2, 4, 6) generated in 2 ms (500 Hz) were coherently added to produce a high-quality compound image. 250 compound images were concatenated into a single Doppler image in 0.5 s. Data were processed using MATLAB. Motion artifacts were corrected using rigid body registration. Power Doppler signal intensity was computed to estimate cerebral blood volume changes. The images were registered to the Allen Mouse Brain Atlas for anatomical localization.

### Immunohistochemistry

Mice were deeply anesthetized and transcardially perfused with either 0.15 M or 2 M NaCl, followed by fixation with 4% paraformaldehyde (PFA) dissolved in 0.15 M or 2 M NaCl. Brains were extracted, post-fixed in 4% PFA overnight at 4□. Brain sections were prepared in 0.15 M or 2 M NaCl using a vibratome (Leica VT1200 S) and treated with 0.3% Triton X-100 for 20 min at room temperature. Then, the free-floating sections were washed with 0.15 M or 2 M NaCl for three times and stained with DAPI. The sections were mounted and imaged with a confocal microscope (Leica Stellaris 5).

### Statistical analysis

Data analysis was carried out in Excel, GraphPad Prism, MATLAB, and Python. All summarized data were presented as the mean ± standard error of the mean (SEM). Statistical significance values between isotonic (0.15 M NaCl) and hypertonic (2 M NaCl) groups were analyzed by two-way ANOVA and denoted as ns (not sgnificant), * (p < 0.05), ** (p < 0.01), or *** (p < 0.001).

## Funding

This work is supported by the following grants:

National Natural Science Foundation of China (32371151)

Guangdong High Level Innovation Research Institute (2021B0909050004) China

Postdoctoral Science Foundation (2024M750595)

China National Postdoctoral Program for Innovative Talents (BX20240093)

## Author contributions

Conceptualization: ZQ.

Methodology: TZ, CJ, JY, HZ, XL, WH, LZ, and ZC Investigation: TZ and CJ

Visualization: TZ and CJ Funding acquisition: ZQ and CJ Project administration: TZ and CJ Supervision: ZQ and HR

Writing – original draft: CJ and ZQ Writing – review & editing: CJ and ZQ

## Data, code, and materials availability

All data are present in the main text or the supplementary materials. The experimental materials and code for data analysis are available from the corresponding author upon reasonable request.

**Fig. S1.**
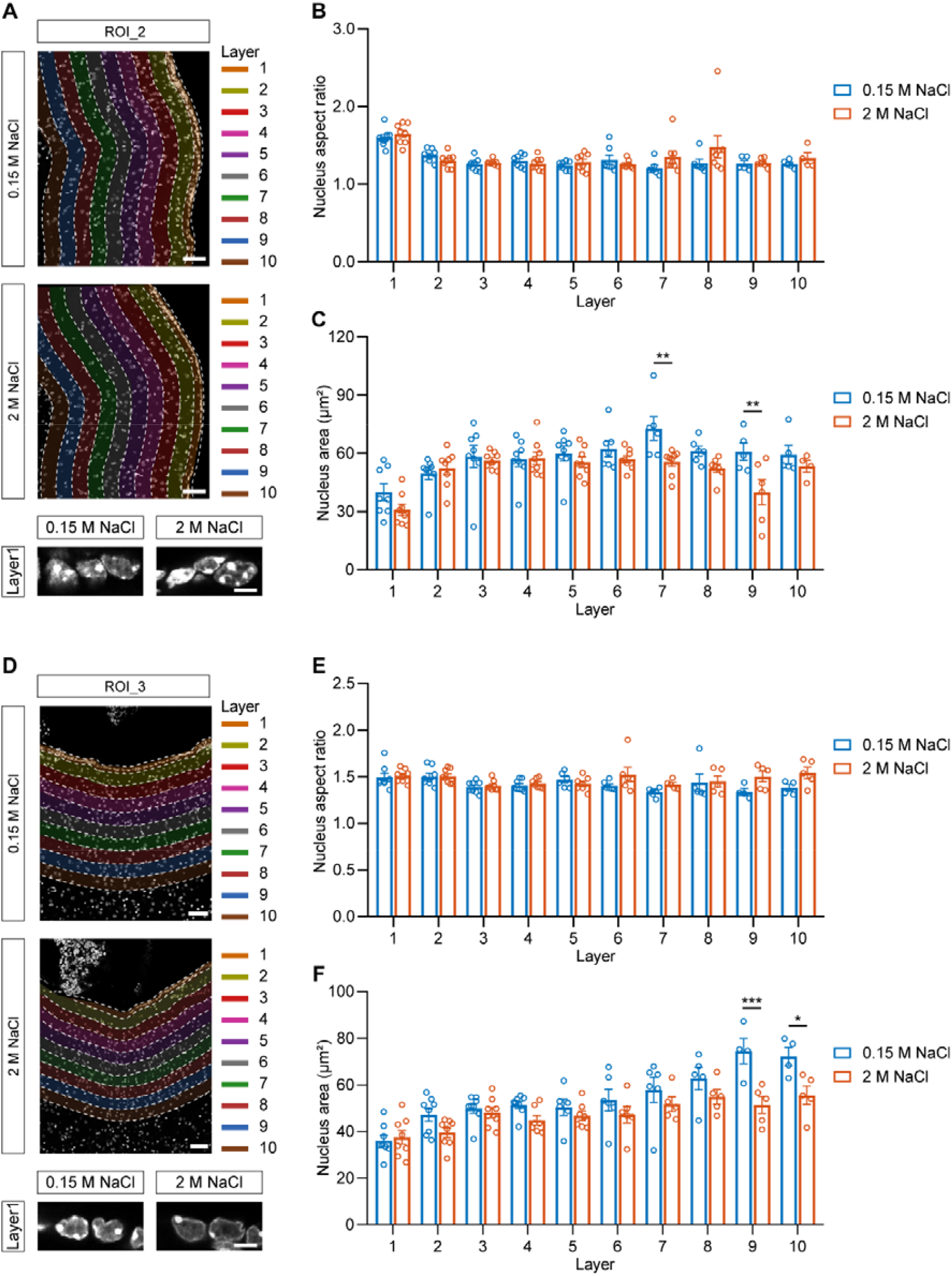
Hypertonicity-induced ventricular expansion deforms periventricular cells. (A) Representative images of the ten layers of nuclei (top and middle, scale bar: 50 µm) and single nucleus in layer 1 (bottom, scale bar: 5 µm) of ROI_2 under isotonic (0.15 M NaCl) or hypertonic (2 M NaCl) condition. (B) Statistical analysis of nucleus aspect ratio of cells in each layer under isotonic (0.15 M NaCl) or hypertonic (2 M NaCl) condition (n = 4-8 mice). (C) Statistical analysis of nucleus area of cells in each layer under isotonic (0.15 M NaCl) or hypertonic (2 M NaCl) condition (n = 4-8 mice). (D) Representative images of the ten layers of nuclei (top and middle, scale bar: 50 µm) and single nucleus in layer 1 (bottom, scale bar: 5 µm) of ROI_3 under isotonic (0.15 M NaCl) or hypertonic (2 M NaCl) condition. (E) Statistical analysis of nucleus aspect ratio of cells in each layer under isotonic (0.15 M NaCl) or hypertonic (2 M NaCl) condition (n = 4-8 mice). (F) Statistical analysis of nucleus area of cells in each layer under isotonic (0.15 M NaCl) or hypertonic (2 M NaCl) condition (n = 4-8 mice). All data were presented as mean ± SEM. Significant differences between isotonic (0.15 M NaCl) and hypertonic (2 M NaCl) groups were analyzed by two-way ANOVA and denoted as * (p < 0.05), ** (p < 0.01), or *** (p < 0.001).

